# Virtual motivation: The psychological and transfer of learning effects of immersive virtual reality practice

**DOI:** 10.1101/2022.12.29.522235

**Authors:** Logan T. Markwell, Joei R. Velten, Julie A. Partridge, Jared M. Porter

## Abstract

Previous research has shown practice within an immersive virtual reality (VR) environment improves real-world (RW) performance. Increased user motivation is one possible advantage of practicing in VR. One recent study showed that an enriched gaming environment led to higher levels of engagement, resulting in a direct learning benefit. The purpose of this study was to compare the intrinsic motivation, engagement, and transfer of learning differences between VR practice and RW practice of the same motor skill. Participants (*n* = 61) were randomly assigned to a RW practice group (*n* = 30) or a VR practice group (*n* = 31) in which they performed a golf putting task. Analyses showed VR practice led to a significantly greater increase in average IMI score than RW practice. Analyses for performance showed there was a significant (*p* < .001) improvement in accuracy (i.e., radial error) from pre to posttest, but the two groups did not differ from one another. Overall, these results partially support our hypotheses and suggest that VR practice led to a greater increase in motivation compared to RW practice. Additionally, these results suggest that VR practice was similarly effective at improving accuracy compared to RW practice. Future research directions are discussed.

## Introduction

Developing a level of motor skill mastery requires a significant amount of quality practice (Ericsson et al., 1993). However, depending on the skill, numerous barriers exist that make it challenging to attain an adequate amount of practice in the pursuit of mastery. For example, many sports require a field or gymnasium and at least one, if not multiple individuals to assist in practice. Pilots require access to planes, helicopters, or simulators to gain experience, which comes at the cost of personal injury or financial expense. The use of virtual reality (VR) practice has been considered a method for overcoming such logistical, inconvenient, and costly obstacles (Michalski et al., 2019a). Professions such as firefighting (Stansfield et al., 2000), surgery (Seymour et al., 2002), aviation (Hays et al., 1992), and sport (Gray, 2017) have used this technology for its assumed benefits. Additionally, VR allows for the utilization and implementation of optimal learning principles that have been rigorously tested for numerous decades (Weiss et al., 2014; Wulf, 2007). Previous research has demonstrated that practice in VR can outperform traditional practice when the VR practice difficulty is adapted based on the individual’s skill level (Gray, 2017). While it is unlikely that VR will completely replace traditional practice, practicing in VR offers numerous advantages, particularly in situations where traditional practice is not available due to cost, safety, and logistics.

Though the research is still in its infancy, empirical evidence supports the conclusion that practicing a motor skill in VR can effectively improve real-world (RW) performance (Harris et al., 2020; Michalski et al., 2019b; Oagaz et al., 2021). This evidence exists for both immersive (e.g., head mount display) and non-immersive (e.g., CAVE system) VR (Gray, 2017; Oagaz et al., 2021). An immersive VR environment is created when a person wears a head mounted system which completely visually encompasses the individual’s field of view into the virtual space. In contrast, a non-immersive VR experience is created when an image is projected onto surrounding walls. A person is able to see the virtual world and the physical world while utilizing a non-immersive VR system. It is important to note that research using immersive VR to investigate changes in motor performance and learning is scarce. Moreover, not all the experiments that have tested the transfer of learning from an immersive virtual environment to a RW environment have provided evidence to support this conclusion (Drew et al., 2020; Harris et al., 2020, experiment 1). For example, Drew et al. (2020) showed that immersive VR practice decreased performance compared to the pretest. Thus, given the minimal amount of research that has examined the transfer of learning in immersive VR, additional research is warranted to understand what does and does not result in RW performance improvements.

The theory of identical elements (Thorndike, 1914) and the transfer-appropriate processing theory (Lee, 1988), which later developed into the identical production model (Singley & Anderson, 1989), have been proposed to explain transfer of learning. These theories purport that similarities must exist between the practice environment and the test environment to achieve a positive transfer of learning effect. More specifically, the transfer appropriate processing theory predicts positive transfer results from *cognitive* processing similarities between the practice and testing environments (Lee, 1988). Whereas the identical elements theory suggests that transfer is due to the similarities between *movement* characteristics executed during practice and testing environments (Thorndike, 1914). Based on research that has tested these theories, it is likely that both, at least to a degree, contribute to a positive transfer of learning effect (Lee, 1988; Singley & Anderson, 1989). Therefore, it can be anticipated that the extent to which positive transfer occurs is influenced by the characteristics of the practiced skill, environmental context, and the cognitive processes shared between the practice and testing environments. Such explanations also align with practice specificity research indicating that transfer of learning will be the greatest when the sources of information available during practice are similar to the information available during post-testing (Proteau, 1992; Proteau et al., 1992). Thus, if the VR environment provides task, environment, and cognitive similarities to those in the physical environment, it is hypothesized that VR practice would facilitate a positive transfer of learning resulting in RW performance improvements.

In addition to the possible RW physical performance improvements following VR practice, VR potentially enhances motivation and engagement when compared to traditional RW practice (Gray, 2019; Lohse et al., 2016). Wulf and Lewthwaite (2016) suggest that psychological properties, such as motivation, are factors that likely contribute to motor performance and learning. Both extrinsic and intrinsic motivation has been shown to benefit learning (Abe et al., 2011; Gruber et al., 2014). Such benefits have been proposed to be indirect through an increased amount of practice (O’Brien & Toms, 2008), and direct via neurophysiological evidence (Anderson et al., 1994; Kempermann et al., 1997). For both intrinsic and extrinsic motivation, neuroimaging studies suggest that the dopaminergic pathways and hippocampus activation facilitate this learning benefit during practice (Adcock et al., 2006; Gruber et al., 2014). Additionally, research in rodents has demonstrated that enriched environments can increase the number of neuron synapses (Anderson et al., 1994) and neuron retention (Kempermann et al., 1997) compared to a non-enriched environment. More recently, work by Lohse et al. (2016) found support for a direct influence on learning in humans. This study showed that a task performed in an enriched gaming environment led to increased engagement and learning compared to the same task performed in a non-enriched environment (Lohse et al., 2016). Thus, a motor skill practiced in VR could elicit similar motivational and engagement improvements compared to an enriched environment leading to possible learning benefits.

To our knowledge, no study has directly assessed whether a motor task performed in immersive VR influences intrinsic motivation or engagement during practice. Thus, the purposes of this study were: 1) to replicate previous studies that have tested the transfer of learning effects following immersive VR motor skill practice and 2) to compare intrinsic motivation and engagement between motor skill practice in an immersive VR environment and RW environment. Based on previous research examining VR transfer of motor learning (Harris et al., 2020; Michalski et al., 2019), we predicted that VR practice would result in performance improvements. Specifically, it was hypothesized that the VR and RW posttest would reveal a significant decrease in radial error (RE) and bivariate variable error (BVE) compared to the pretest. Moreover, it was predicted that no significant group differences would be found for RE or BVE. Furthermore, it was predicted that the VR posttest would not reveal significant group differences. Lastly, we predicted that VR practice would lead to greater intrinsic motivation and engagement following practice compared to RW practice.

## Method

### Participants

A total of 64 university students (males = 21; females = 43) between the ages of 18-30 (*M* = 21.97, *SD* = 2.45) volunteered to participate in the present experiment. An a priori power analysis was performed using G*Power 3.1.9 (Faul et al., 2007; Faul et al., 2009). Based on the effect size from pilot data (*f* = .190), an alpha = .05, and power = .80, the projected sample size needed was approximately *N* = 58 for a within-between subject comparison. Of the 64 students which volunteered for the study, a total of 61 participants completed the experiment. Participants were informed they would practice a golf putting task and that they would use VR but were naïve to the purpose of the study. The university’s institutional review board approved the study, and the students completed an informed consent form prior to participation.

### Task and Apparatus

Data collected for this experiment took place in a climate-controlled research laboratory. The RW golf putting task was performed on an artificial grass carpeted surface (1.829 × 3.658 m) inside the laboratory. Participants used a standard length (90 cm) golf putter to putt a regulation-sized golf ball towards a target the size of a standard golf cup hole (diameter 10.795 cm). The hole was 2.438 meters away from the starting line. A web camera was fixed perpendicularly above the target to capture the golf ball’s position relative to the center of the target. The camera application (Microsoft Corporation; version 2021.105.10.0) on an Alienware computer was used for video capture. Tracker software (version 6.0.1) was used to determine the x and y coordinates of the at-rest golf ball position.

The Oculus Quest 2 VR headset and Cloudlands VR Minigolf application were used to create a virtual miniature golf putting course that was designed to replicate the course in the physical environment. The exact shape and dimensions of the RW putting surface were replicated in the VR environment. Participants used a virtual golf putter to putt a virtual ball into a virtual hole while wearing a Oculus Quest 2 headset and holding one Oculus controller in their dominant hand. The researcher recorded the golf ball’s Euclidian distance from the hole, provided by the Cloudlands VR Minigolf application.

### Procedure

Participants were randomly assigned into one of two groups: VR practice (*n* = 31) or RW practice (*n* = 30). After signing the consent form, participants completed the mini simulator sickness questionnaire (MSSQ) to evaluate their potential risk of motion sickness. Participants that scored 26 or greater on the MSSQ were excluded from the study. Following the MSSQ, the participants completed an intrinsic motivation inventory (IMI). After the participants completed the questionnaires, the researcher provided instructions followed by a demonstration of the golf putting task. Participants were instructed to hit the ball to the center of the target with the goal of making the golf ball come to rest in the center of the target or as close to the center of the target as possible.

The experiment took place over two consecutive days. On day one, participants completed a pre-practice IMI, a golf putting pretest, golf putting acquisition phase in VR or RW, the O’Brien engagement scale, and a second IMI post-practice. The questionnaires were counterbalanced across participants to control for a possible order effect. The posttest occurred on day two. The pre- and posttest phases were identical for both groups (i.e., VR & RW). During the pre- and post-testing phases, participants putted a golf ball ten times on the carpeted surface towards the center of the target. The practice phase consisted of 50 total putts within the respective environment (i.e., VR or RW). Participants returned within 48 hours to complete the posttests. The posttests included 10 golf putts in the RW environment and 10 golf putts in the VR environment. The two posttests were counterbalanced across participants. Please see figure 1 for a schematic representation of the experimental design.

**Figure 1.**
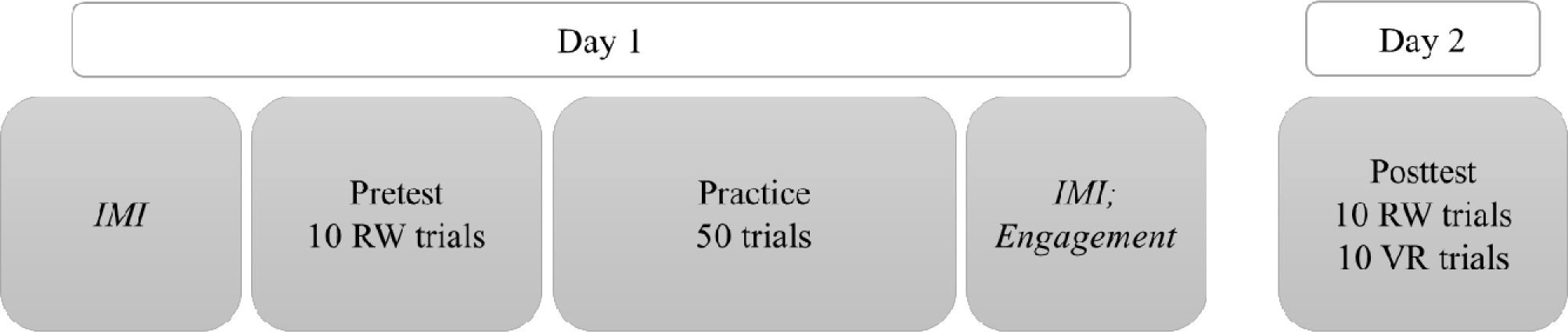
Experiment procedure.

Putting performance data were collected to determine participants’ accuracy (i.e., radial error) and precision (i.e., bivariate variable error). The putting target was considered the origin of a two-dimensional grid with the coordinates 0,0. Radial error (RE), a two-dimensional equivalent of absolute error, was calculated for each trial using the Pythagorean theorem to calculate the Euclidian distance of the two closest points between the golf ball and the center of the target.

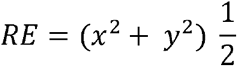

Bivariate variable error (BVE), the two-dimensional equivalent of variable error, was calculated by taking the square root of the squared mean distance of each trial from the centroid (*c*) of each block of *k* trials.

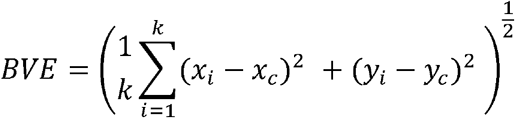

### Data Analysis

Data were analyzed using IBM SPSS Statistics version 28.0.0. There was seven separate 2 (group) x 2 (test) repeated measures analysis of variance (ANOVAs) used to assess the average intrinsic motivation scores and four subscales (interest/enjoyment, perceived competence, effort/importance, value/usefulness), accuracy (radial error), and precision (bivariate variable error). An independent samples t-test was used to determine RE group differences in the VR posttest. Lastly, an independent samples t-test was used to determine engagement score differences between groups.

## Results

### Performance Variables

#### Accuracy - Radial Error (RE)

A 2 (group) x 2 (test) repeated measures ANOVA was used to determine accuracy differences between groups and tests. The analysis revealed a main effect for test, *F*(1, 60) = 15.674, *p <* .001, *η*_*p*_^*2*^ = .207. Pairwise comparisons for the test indicated that both groups significantly decreased radial error from pretest to posttest, *p* < .001. No significant differences were observed for the test x group, *p* = .132, or between-group effects tests, *p* = .738 (see figure 2). Additionally, to assess group differences during the VR posttest, an independent samples t-test revealed no RE group differences between RW (*M =* .765) and VR (*M =* .719), *t*(58) = .763, *p =* .449.

**Figure 2.**
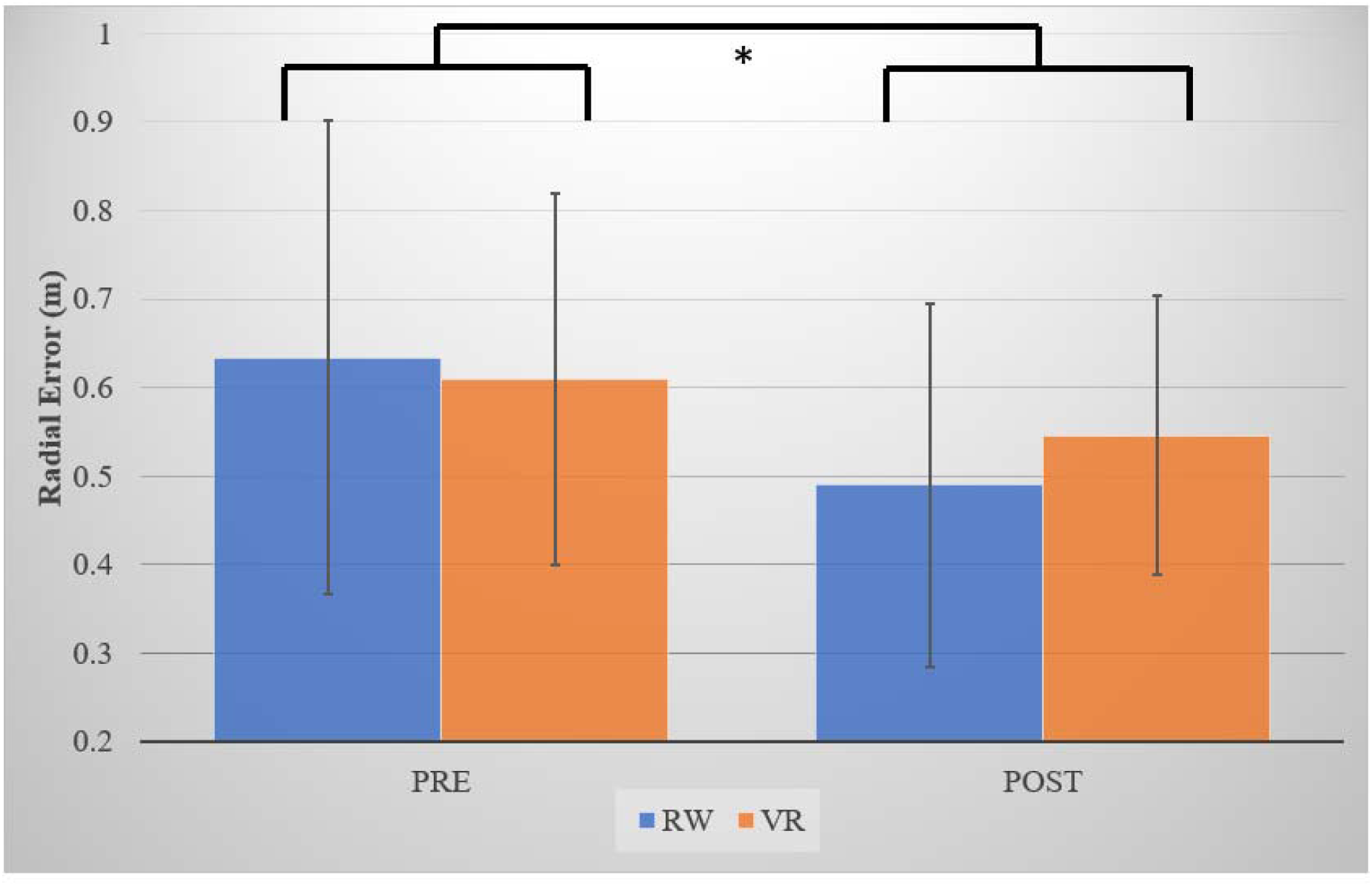
Mean radial error (RE) for pre- and posttests for virtual reality (VR) and real-world (RW) practice groups. The * indicates significant differences. The error bars represent standard deviation (SD).

#### Precision (Bivariate Variable Error)

A 2 (test) x 2 (group) repeated measures ANOVA was used to determine precision differences between tests and groups. There was no significant main effect for the test *F*(1, 60) = 2.536, *p =* .117, *η*_*p*_^*2*^ = .041, the test x group interaction *F*(1, 60) = .025, *p =* .874, *η*_*p*_^*2*^ = .000, or the between-group effects tests *F*(1, 60) = 1.579, *p =* .214, *η*_*p*_^*2*^ = .026.

### Psychological Variables

#### Intrinsic Motivation Inventory

##### Average Intrinsic Motivation

A 2 (group) x 2 (test) repeated measures ANOVA was used to determine average intrinsic motivation score differences between the two groups and the two tests. The analysis revealed a significant main effect for test *F*(1, 59) = 37.827, *p <* .001, *η*_*p*_^*2*^ = .391. The analysis also revealed a significant test x group interaction, *F*(1, 59) = 8.379, *p =* .005, *η*_*p*_^*2*^ = .124. Furthermore, the test of between-group effects revealed a non-significant effect, *p* = .173. Considering the significant interaction, pairwise comparisons were made (see figure 3). Pairwise comparisons revealed no significant differences between the VR (*M =* 4.46, *SD =* 0.76) and RW (*M =* 4.36, *SD =* 0.83) groups at pretest, *p* = .635, but showed that the VR scores (*M =* 5.05, *SD =* 0.86) were significantly higher during the posttest compared to RW scores (*M =* 4.57, *SD =* 0.94), *p =* .045. Additional pairwise comparisons showed that both the VR, *p <* .001, and RW, *p =* .026, scores significantly increased from pre-to post-test. IMI scores are represented in Figure 3.

**Figure 3.**
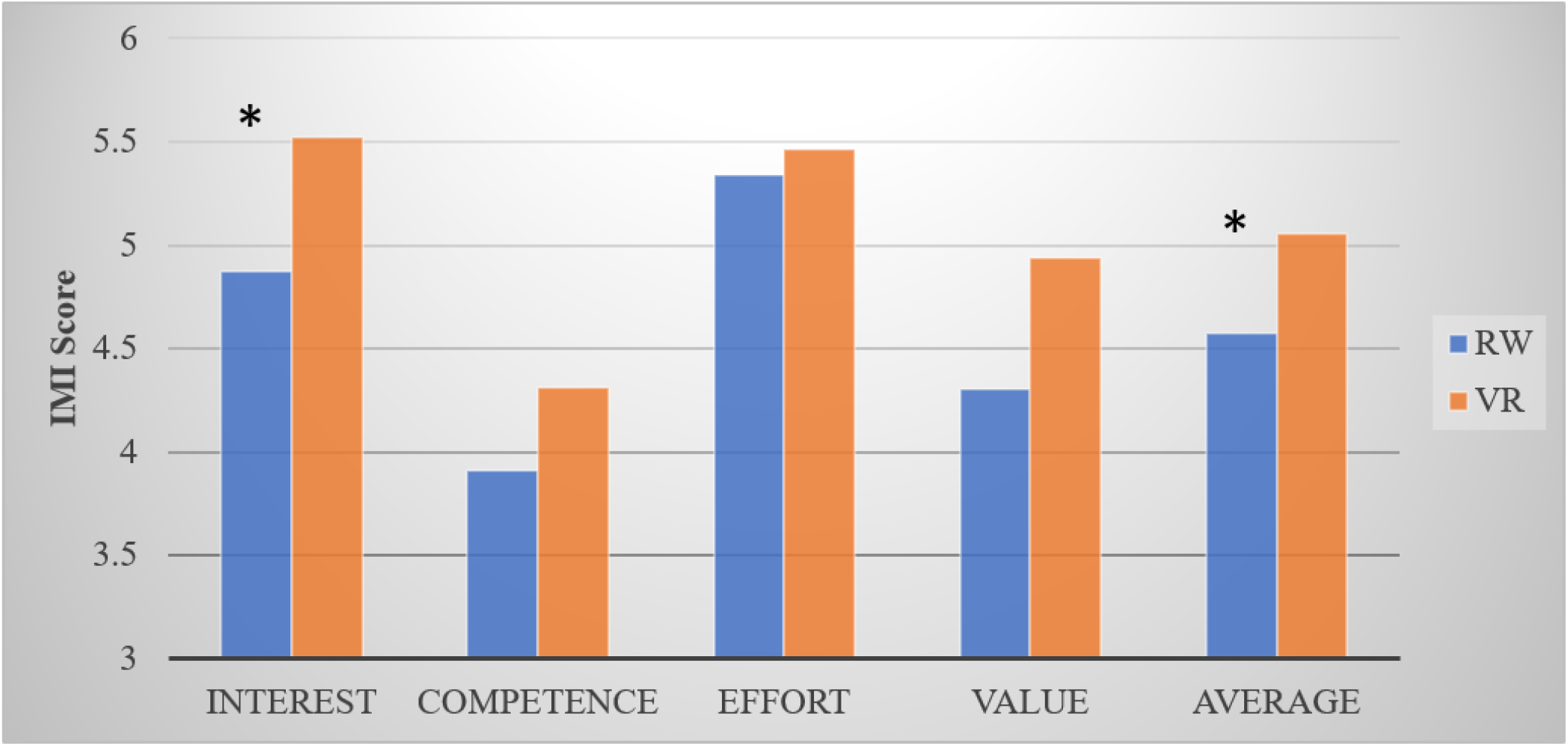
Intrinsic Motivation Inventory Score Differences. The * indicates significant differences between groups.

##### Interest/Enjoyment

A 2 (group) x (test) repeated measures ANOVA was used to determine interest/enjoyment score differences between the two groups during the pre- and post-test. The analysis showed a significant main effect for test *F*(1, 59) = 54.394, *p <* .001, *η*_*p*_^*2*^ = .480. The analysis also found a significant test x group interaction *F*(1, 59) = 11.795, *p =* .001, *η*_*p*_^*2*^= .167. The test of between-group effects was non-significant, *p =* .213. Pairwise comparisons showed that the VR (*M* = 4.50, *SD* = .188) and RW (*M* = 4.50, *SD* = .191) scores were not significantly different, *p =* .993. However, the comparison revealed the posttest VR score (*M* = 5.521, *SD* = .195) was significantly higher compared to the RW score (*M* = 4.87 *SD* = .199). Additional pairwise comparisons showed that the RW, *p =* .008, and VR, *p <* .*001*, groups significantly increased scores from pre-to post-test.

##### Perceived Competence

A 2 (group) x 2 (test) repeated measures ANOVA revealed a significant main effect for test *F*(1, 59) = 10.799, *p =* .002, *η*_*p*_^*2*^ = .155. The competence scores for both groups significantly increased from pre-to posttest. No significant interaction, *p* = .147, or between-group effects, *p =* .447, were found.

##### Effort/Importance

To test for group and test differences for effort/importance scores, a 2 (group) x 2 (test) repeated measures ANOVA was used. No significant main effects were found for test, *p* = .254, or between groups, *p* = .765.

##### Value/Usefulness

A 2 (group) x (test) repeated measures ANOVA was used to determine value/interest score differences between groups and tests. The analysis revealed a significant main effect for test *F*(1, 59) = 24.452, *p <* .001, *η*_*p*_^*2*^ = .293. The test x group interaction did not reveal a significant main effect, *p* = .105. Similarly, the between-group main effect did not yield a significant effect, *p* = .056. Pairwise comparisons for test showed that both groups significantly increased scores from pre-to posttest.

#### Engagement

An independent samples t-test was used to compare posttest engagement scores between groups. The t-test revealed that engagement scores for the VR group (*M =* 4.35, *SD =* .668) were not significantly different to the RW group (*M =* 4.362, *SD =* .467), *t*(58) = .032, *p* = .487.

## Discussion

The present study examined the effects of immersive VR practice on the transfer of learning of a golf-putting task. Additionally, this study investigated the effects of VR practice on intrinsic motivation and engagement. It was predicted that VR practice would facilitate a positive transfer of learning to a RW environment, and the performance improvements would be similar between the VR and RW groups. It was also hypothesized that VR practice would lead to higher levels of intrinsic motivation and engagement than RW practice. The results partially supported our hypotheses. A detailed explanation of our results and avenues for future research are discussed below.

### Performance Variables

The performance measurement predictions were partially supported. Specifically, the analysis showed that the RE in both groups significantly decreased from the pretest to the posttest, and no differences were observed between groups. These results suggest that immersive VR practice led to a positive transfer of learning and resulted in a motor learning effect that was relatively similar compared to RW practice. In other words, both forms of practice (i.e., VR and RW) resulted in similar golf putting enhancements. These findings are congruent with previous research and support the conclusion that VR practice results in RW motor performance improvement (Harris et al., 2020, experiment 2; Michalski et al., 2019b; Oagaz et al., 2021). For example, Michalski et al. (2019b) found that VR practice led to significant performance improvements in table tennis strokes and greater performance compared to a no-practice control group. Similarly, a second experiment by Harris et al. (2020) showed that both RW and VR practice of a golf putting task led to increased RW accuracy, and no differences were found between practice conditions.

Although previous research has demonstrated that practicing a motor skill in virtual reality improves the performance of the same skill in the real world, some studies have found conflicting results. For example, two recent experiments investigating this topic contradict the findings reported in the present study. These studies demonstrated that VR practice degraded RW motor performance (Drew et al., 2020; Harris et al., 2020, experiment 1). Unlike the present study, and previous research supporting a positive transfer of learning effect (Michalski et al., 2019; Oagaz et al., 2021), the experiments that found performance impairments following VR practice performed a posttest immediately after practice rather than at least one day following acquisition (Drew et al., 2020; Harris et al., 2020, experiment 1). It is important to note this methodological difference between the present study and earlier experiments (Drew et al., 2020; Harris et al., 2020, experiment 1) as it is well accepted in the skill acquisition literature that the effective assessment of motor learning should include a minimum of one day between the conclusion of practice and the assessment of learning (Schmidt et al., 2019).Thus, it is possible that the amount of time between VR practice and the RW posttest could influence the transfer of learning. This difference in methodology may help explain why our results differed from those reported by Drew et al. (2020) and Harris et al. (2020, experiment 1)

While other VR studies (e.g., Harris et al., 2020; Michalski et al., 2019) have measured performance accuracy, none have used BVE as a measure of practice performance or motor learning. Moreover, non-VR research has commonly used BVE to assess motor precision (e.g., Frank et al., 2016; Hancock et al., 1995). Additionally, BVE has been suggested to be an essential measure of motor learning (Schmidt et al., 2019). The analysis of BVE in the current study did not support the experimental hypothesis. Contrary to the hypothesis, the results reported here suggest that neither group increased golf putting precision as a result of practice. It is worth noting that previous research has shown that precision and accuracy can be influenced differently by practice (Kumar et al., 2017). Nonetheless, improvements in precision for both groups were predicted. During the acquisition phase of this experiment, the amount of variability between each putting trial was minimal. That is, participants initiated all 50 trials from the same start location. A large amount of research has shown a significant motor learning advantage when variability between trial start locations is introduced (Shea & Kohl, 1990). Thus, it is possible that the lack of variability between practice trials negated the practice effect of increased precision. Future research comparing the transfer of learning between VR and RW practice should use a practice schedule that induces practice variability. Additionally, this future line of research will reveal if introducing a learner to variability through VR is equally effective to experiencing practice variability in the real-world. This is important to understand from both a theoretical and practical point of view.

Lastly, the results of the RE analysis during the VR posttest revealed no group differences. Contrary to the present study’s results, Harris et al. (2020) showed that only the VR group improved in the VR environment. However, unlike Harris et al. (2020), the present study only assessed posttest group differences rather than pre-to posttest changes. Harris et al. (2020) did not find group differences but only a significant improvement in the VR group. Therefore, future research could include a VR pretest to determine changes over time.

### Psychological Variables

Regarding the psychological measurements, the experimental hypotheses were partially supported in that the results from the intrinsic motivation measurements support the current study’s predictions. Specifically, it was shown that VR and RW practice significantly increased participants’ average intrinsic motivation. Moreover, VR led to a greater increase in intrinsic motivation compared to RW practice, as evidenced by the significantly higher posttest scores. When examined by the subscales, both VR and RW practice significantly increased interest and enjoyment. However, practice in VR led to greater increases in both measures. The results for perceived competence, effort, and importance showed that both VR and RW groups increased scores, whereas there was no change observed in value and usefulness for either group.

The results of the present study confirm the prediction made by Gray (2019), suggesting that practicing a motor task in VR creates a learning environment that is more motivating compared to RW practice. Such a finding is valuable given that increasing intrinsic motivation has been shown to facilitate motor performance (Wulf & Lewthwaite, 2016), and previous work has demonstrated indirect (Hunicke et al., 2004) and direct (Abe et al., 2011; Gruber et al., 2014) learning benefits as a product of elevated motivation. It is worth noting that the increase in intrinsic motivation occurred following only one practice session. Thus, whether this motivational increase remains elevated after multiple practice sessions in VR is unknown. Interestingly, previous research investigating the novelty of learning environments has shown that when individuals are exposed to a new learning context, perceived novelty increases, which is associated with an increase in intrinsic motivation (Jeno et al., 2019). Additionally, the appraisal of novelty has been shown to predict higher levels of interest (Adachi et al., 2017). However, the increase in novelty and motivation has been shown to decrease with repeated exposure to the learning context (Keller & Suzuki, 2004). Such findings may explain the intrinsic motivation results found in the current study. That is, the increase in intrinsic motivation could be a product of VR creating a novel learning environment. This is further supported in that the only subscale that revealed group differences during the posttest was interest and enjoyment, consistent with findings reported in previous novelty research (Adachi et al., 2017). Therefore, further investigation is warranted to understand if this observed increase in motivation remains elevated after repeated exposure to VR practice.

Contrary to the motivation hypothesis, the prediction made for elevated engagement was not supported by the findings of this experiment. Unlike Lohse et al. (2016), the results of this study revealed no significant differences in engagement between the VR and RW groups during the posttest. The methodology in the experiment by Lohse et al. (2016) compared a computer task performed within a RW sterile environment compared to the same task being practiced in a dynamic gamified virtual environment. In the present study, the environments were different, but it is possible the environments were not different enough (e.g., sterile vs. gamified) to result in self-reported engagement differences. For example, the VR environment in the present study was not gamified, unlike the environment utilized by Lohse et al. (2016). Therefore, it is possible that this lack of gamification contributed to the lack of observed differences. Thus, additional research is warranted to understand whether there are engagement differences between VR and RW practice as a result of changing the dynamics of both environments.

### Limitations and Conclusion

The present study shows that VR practice results in similar RW performance improvements and higher motivation and interest compared to RW practice. These results are promising for the future development of VR technology as few studies have investigated the transferability of VR practice of motor skills being performed in a RW environment. This study is one of the few to provide evidence for a positive transfer of learning effect, extending the findings reported in previous research (Harris et al., 2020, experiment 2; Michalski et al., 2019; Oagaz et al., 2021). Moreover, this is the first study to compare intrinsic motivation and engagement differences during motor skill practice between VR and RW environments. However, it is worth noting that the amount of practice performed is a primary limitation. The practice specificity hypothesis purports two primary claims: 1) if information differences exist between the acquisition and testing phases, there will likely be a decrease in performance during the testing phase, and 2) this negative performance effect will increase as the amount of practice increases (Proteau, 1992). Thus, given that information differences likely exist between a VR and RW environment, it is predicted that an extended amount of VR practice would lead to transfer of learning differences compared to RW practice. Such performance differences should be observed between VR and RW practice during a RW posttest. Additionally, the increase in intrinsic motivation may simply be a result of novelty during VR practice (Jeno et al., 2019). If such an observation is due to novelty, increasing the amount of VR practice might decrease novelty and thereby decrease motivation (Adachi et al., 2017). Thus, increasing the amount of VR practice and measuring performance and intrinsic motivation across multiple time points is one way to overcome such methodological limitations.

## References

Abe, M., Schambra, H., Wassermanm, E. M., Luckenbaugh, D., Schweighofer, N., & Cohen, L. G. (2011). Reward improves long-term retention of a motor memory through induction of offline memory gains. Current Biology, 21, 557–562.

Adachi, P. J., Ryan, R., Frye, J., McClung, D., & Rigby, C. (2017). “I can’t wait for the next episode!” investigating the motivational pull of television dramas through the lens of self-determination theory. Motivation Science, 4, 78–94.

Adcock, R. A., Thangavel, A., Whitfield-Gabrieli, S., Knutson, B., & Gabrieli, J. D. (2006). Reward-motivated learning: Mesolimbic activation precedes memory formation. Neuron, 50, 507–517.

Anderson, B. J., Li, X., Alcantara, A. A., Isaacs, K. R., Black, J. E., & Greenouch, W. T. (1994). Glial hypertrophy is associated with synaptogenesis following motor-skill learning, but not with angiogenesis following exercise. Glia, 11, 73–80.

Drew, S. A., Awad, M. F., Armendariz, J. A., Gabay, B., Lachica, I. J., & Hinkel-Lipsker, J. W. (2020). The trade-off of virtual reality training for dart throwing: A facilitation of perceptual-motor learning with a detriment to performance. Frontiers in Sports and Active Living.

Ericsson, K. A., Krampe, R. T., & Tesch-Romer, C. (1993). The role of deliberate practice in the acquisition of expert performance. Psychological Review, 100, 363–406.

Faul, F., Erdfelder, E., Buchner, A., & Lang, A. G. (2009). Statistical power analyses using G*Power 3.1: Tests for correlation and regression analyses. Behavior Research Methods, 41, 1149–1160.

Faul, F., Erdfelder, E., Lang, A., & Buchner, A. (2007). G*Power 3: A flexible statistical power analysis program for the social, behavioral, and biomedical sciences. Behavior Research Methods, 39, 175–191.

Frank, C., Land, W. M., & Schack, T. (2016). Perceptual-cognitive changes during motor learning: The influence of mental and physical practice on mental representation, gaze behavior, and performance of a complex action task. Frontiers in Psychology, 6, 1981.

Gray, R. (2017). Transfer of training from virtual to real baseball batting. Frontiers in Psychology, 8, 2183.

Gray, R. (2019). Virtual environments and their role in developing perceptual-cognitive skills in sports. In A. M. Williams & R. C. Jackson (Eds.). Anticipation and decision making in sport (pp. 342–358). Routledge.

Gruber, M. J., Gelman, B. D., & Ranganath, C. (2014). States of curiosity modulate hippocampus-dependent learning via the dopaminergic circuit. Neuron, 22, 486–496.

Hancock, G. R., Butler, M. S., & Fischman, M. G. (1995). On the problem of two-dimensional error scores: measures of analyses of accuracy, bias, and consistency. Journal of Motor Behavior, 27, 241–250.

Harris, D. J., Buckingham, G., Wilson, M. R., Brookes, J., Mushtaq, F., Mon-Williams, M., & Vine, S. J. (2020). The effect of virtual reality environment on gaze behavior and motor skill learning. Psychology of Sport and Exercise, 50, 101721.

Hays, R. T., Jacobs, J. W., Prince, C., & Salas, E. (1992). Flight simulator training effectiveness: A meta-analysis. Military Psychology, 4, 63–74.

Hunicke, R. LeBlanc, M., & Zubek, R. (2004). “MDA: formal approach to game design and game research,” in Proceedings of the Challenges in Game AI Workshop, 19th National Conference on Artificial Intelligence, San Jose, CA.

Jeno, L. M. Vandvik, V., Eliassen, S. Grytnes, J. (2019). Testing the novelty effect of an m-learning tool on internalization and achievement: A Self-Determination Theory Approach. Computers & Education, 128, 398–413.

Keller, J. N. & Suzuki, K. (2004). Learner motivation and E-learning designs: A multinationally validated process. Journal of Education Media, 29, 229–239.

Kempermann, G., Kuhn, H. G., & Gage, F. H. (1997). More hippocampal neurons in adult nice living in an enriched environment. Nature, 368, 493–495.

Kumar, A., Tanaka, Y., Grigoriadis, A., Grigoriadis, J., Trulsson, M., & Svensson, P. (2017). Training-induced dynmaics of accuracy and precision in human motor control. Scientific reports, 7, 67984.

Lee, T. D. (1988). Transfer-appropriate processing: A framework for conceptualizing practice effects in motor learning. In: Meijer, O. G, Roth, K. (eds.). Advances in psychology. vol. 50. Amsterdam: North Holland, pp. 201–215.

Lohse, K., Boyd, L., & Hodges, N. (2016). Engaging environments enhance motor skill learning in a computer gaming task. Journal of Motor Behavior, 48, 172–182.

Michalski, S. C., Szpak, A., & Loetscher, T. (2019a). Using virtual environments to improve RW motor skills in sports: A systematic review. Frontiers in Psychology, 10.

Michalski, S.C., Szpak, A., Saredakis, D., Ross, T.J., Billinghurst, M., & Loetscher, T. (2019b). Getting your game on: Using virtual reality to improve real table tennis skills. PLoS ONE, 14, e0222351.

O’Brien, H. L. & Toms, E. (2008). What is user engagement? A conceptual framework for defining user engagement with technology. Journal of the American Society for Information Science and Technology, 59, 938–955.

Oagaz, H., Schoun, B., & Choi, M. (2021). Performance improvement and skill transfer in table tennis through training in virtual reality. IEEE Transactions on visualization and computer graphics.

Proteau, L. (1992). On the specificity of learning and the role of visual information for movement control. In L. Proteau & D. Elliott (Eds.), Vision and motor control (pp. 67–103). North Holland.

Proteau, L., Martenuik, R., & Levesque, L. (1992). A sensorimotor basis for motor learning: Evidence indicating specificity of practice. The Quarterly Journal of Experimental Psychology A: Human Experimental Psychology, 44, 557–575.

Schmidt, R. A., Lee, T. D., Winstein, C., Wulf, G., & Zelaznik, H. K. (2019). Motor Control and Learning: A behavioral emphasis. Human Kinetics.

Seymour, N. E., Gallagher, A. G., Roman, S. A., O’Brien, M. K., Bansal, V. K., Andersen, D. K., & Satava, R. M. (2002). Virtual reality training improves operating room performance: Results of a randomized, double-blinded study. Annals in Surgery, 236, 458–463.

Shea, C. H. & Kohl, R. M. (1990). Specificity and variability of practice. Comparative Study, 61, 169–177.

Singley, M. K., & Anderson, J. R. (1989). The transfer of cognitive skill. arvard University Press.

Stansfield, S., Shawver, D., Sobel, A., & Prasad, M. (2000). Design and implementation of a virtual reality system and its application to training medical first responders. Presence Teleoperators & Virtual Environment, 9, 524–556.

Thorndike E. L. (1914). The psychology of learning. Teachers College.

Weiss, P.L., Keshner, E.A., & Levin, M.F. (2014). In P. Sharkey P (Ed.) Applying Virtual Reality technologies to motor rehabilitation: Virtual reality technologies for health and clinical Applications. Vol. 1 Springer.

Wulf, G. & Lewthwaite, R. (2016). Optimizing performance through intrinsic motivation and attention for learning: The OPTIMAL theory of motor learning. Psychonomic Bulletin & Review, 23, 1383–1414.

Wulf, G. (2007). Attention and motor skill learning. Human Kinetics.

